# Two-stage Dynamic Deregulation of Metabolism Improves Process Robustness & Scalability in Engineered *E. coli*

**DOI:** 10.1101/2020.08.30.274290

**Authors:** Zhixia Ye, Shuai Li, Jennifer N. Hennigan, Juliana Lebeau, Eirik A. Moreb, Jacob Wolf, Michael D. Lynch

## Abstract

We report improved strain and bioprocess robustness as a result of the dynamic deregulation of central metabolism using two-stage dynamic control. Dynamic control is implemented using combinations of CRISPR interference and controlled proteolysis to reduce levels of central metabolic enzymes in the context of a standardized two-stage bioprocesses. Reducing the levels of key enzymes alters metabolite pools resulting in deregulation of the metabolic network. The deregulated network is more robust to environmental conditions improving process robustness, which in turn leads to predictable scalability from high throughput small scale screens to fully instrumented bioreactors as well as to pilot scale production. Additionally, as these two-stage bioprocesses are standardized, a need for traditional process optimization is minimized. Predictive high throughput approaches that translate to larger scales are critical for metabolic engineering programs to truly take advantage of the rapidly increasing throughput and decreasing costs of synthetic biology. In this work we demonstrate that the improved robustness of *E. coli* strains engineered for the improved scalability of the important industrial chemicals alanine, citramalate and xylitol, from microtiter plates to pilot reactors.

## Introduction

Fermentation processes have made rapid advancements in recent years due to technology developments in the fields of fermentation science and synthetic biology, as well as metabolic and enzyme engineering.^1–4^ Despite these substantial advances, most successful examples of rational and directed strain engineering approaches have also greatly relied on numerous and often lengthy cycles of trial and error. ^5–7^ One reason for continued trial and error is a fundamental lack of robustness in many biobased processes when compared to more traditional process chemistry. The metabolic networks of all living systems, including microbes, are highly regulated and respond to environmental conditions.^8,9^ These responses, while highly adaptive, make strain engineering difficult when environmental conditions can often change, even if only subtly, between experimental and process conditions. ^8,10–12^ This lack of biological robustness has two primary consequences, the first is a lack of standardization and the second is difficulty in scaling.

Historically, due to the large impact of environmental variables on bioprocess performance, traditional process development and optimization, has been a core essential function in the development of almost all industrial bioprocesses. In addition to strain modifications, media formulations, feeding and oxygenation strategies (in the case of aerobic processes) as well as various seed protocols are routinely optimized for a given strain and product. A lack of robustness has not only made these workflows critical, but has also resulted in a field where each successful process often represents almost a unique, artisanal solution. Due to the accepted need for significant process development, to many, strain and process standardization is not feasible.

Secondly, the lack of robustness has also made scaling up of bioprocesses a significant challenge and risk. This occurs not only when scaling from screening systems to instrumented reactors but also from lab scale production to the pilot and commercial scales. Consequently, results obtained from high throughput studies or even shake flask experiments often do not translate, even in the same microbe, to a different context, such as a larger scale. ^8,12,13^ This is traditionally one factor making the scale up of fermentation based processes difficult, as small scale screening studies do not always readily translate to larger scale production processes. This is particularly true with aerobic fermentations. Oftentimes specific screening protocols are developed to “scale down” a process, and as a result screening approaches also tend to become artisanal with significant customization based on a given strain, process or product. This has led to the development of specialized, complex micro-reactor systems for scale down studies, where process conditions can be better controlled.^8,14–16^ The efforts in developing these systems again stem from a lack of robustness in biological systems and a concerted effort to improve the throughput of process optimization.

Despite the fact that a lack of robustness is a primary challenge facing metabolic engineering and bioprocess development it is often taken for granted as a necessary challenge to be solved later (upon commercial development) and is not considered as a primary output in a majority of strain engineering or synthetic biology studies. Even the production of recombinant protein, a relatively simple product, can be impacted by a lack of robustness. For example, we have recently reported that improved promoter robustness corresponds to increased scalability from high throughput studies to instrumented bioreactors. ^17^

In order to address the primary challenge of robustness in the more complicated production of chemicals via engineered metabolism, in this study we investigate the use of dynamic metabolic control in two-stage cultures to improve process robustness and scalability. Dynamic metabolic control, which dynamically alters the metabolic network, has become a common strategy in metabolic engineering, with development of numerous tools and applications. ^18–27^ Two-stage dynamic control, which decouples growth from production, offers additional potential benefits in large scale bioprocesses, enabling metabolic states to be pushed beyond the boundaries required for growth. ^8,21,22,24,28^

We have recently implemented two-stage dynamic control in the commonly used and well characterized microbe, *E. coli*, by utilizing synthetic metabolic valves, which rely on the combination of proteolysis and gene silencing to dynamically reduce levels of key enzymes in the context of a standardized two-stage phosphate depleted process (as illustrated in Figure 1) ^17,21,22,29^ Proteolysis is accomplished by appending C-terminal degron (DAS+4) tags to a given gene and silencing via expression of the native *E. coli* CRISPR CASCADE system as well as silencing gRNAs (from pCASCADE plasmids).^30,31^ Importantly, in these studies where we produced the organic acid citramalate ^22^ and the sugar alcohol xylitol, ^21^ we identified that dynamic deregulation of metabolism was a fundamental mechanism to improve stationary phase metabolic fluxes. Specifically, i) reducing citrate synthase (GltA) levels reduced *α*-ketoglutarate pools alleviating *α*-ketoglutarate mediated inhibition of glucose uptake,^22^ ii) reducing glucose-6-phosphate dehydrogenase (Zwf) levels reduces NADPH pools activating the SoxRS regulon and increasing expression of pyruvate ferredoxin oxidoreductase and acetyl-CoA flux,^22^ and iii) decreasing enoyl-ACP reductase (FabI) activity decreases fatty acid metabolite pools alleviating inhibition of the membrane bound transhydrogenase.^21^ The deregulation of central metabolic pathways enabled relatively high titers in instrumented reactors, reaching 125g/L in the case of citramalate and 200g/L in the case of xylitol.

**Figure 1:**
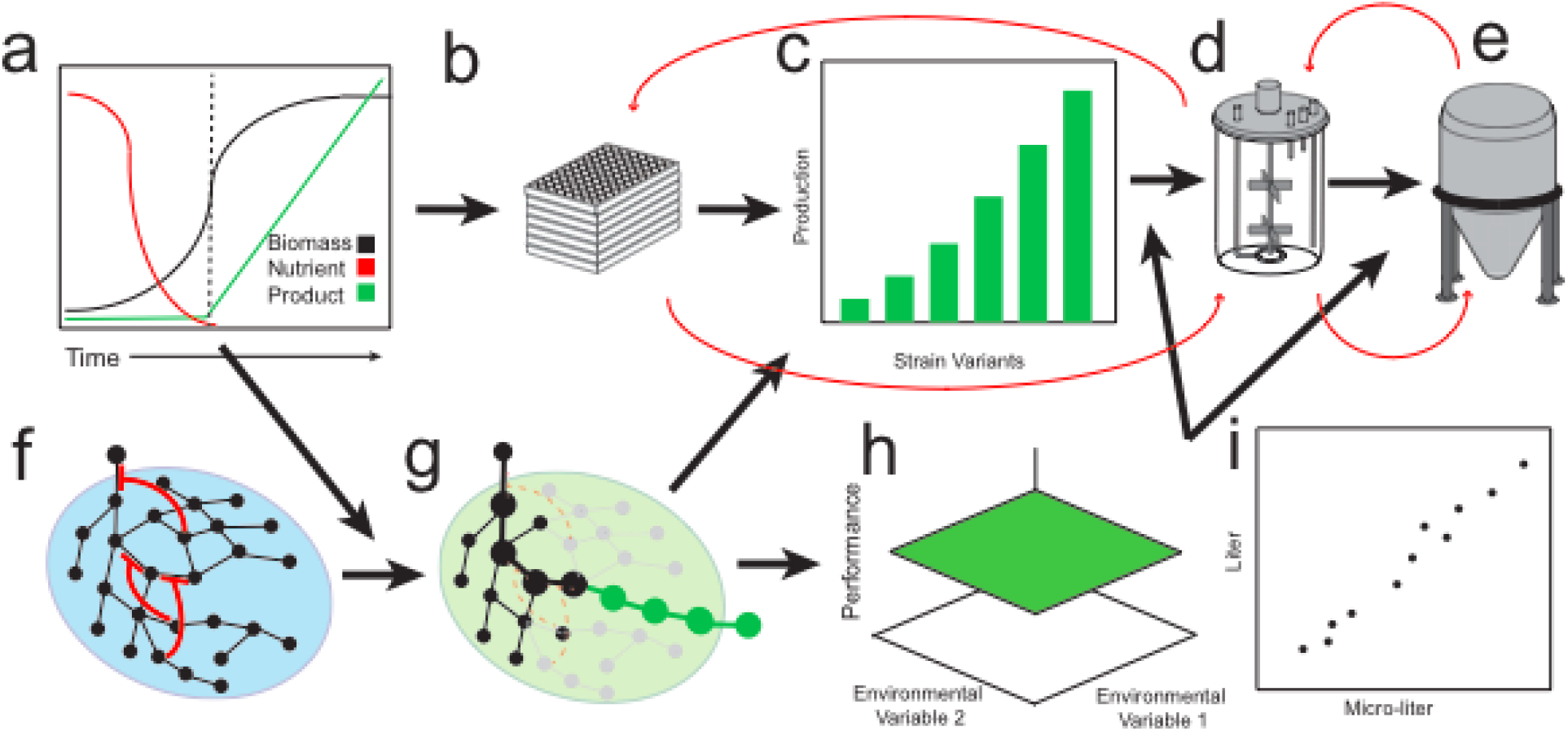
Improved robustness and scalability using two-stage dynamic metabolic control. (a) This is accomplished in a standardized two-stage bioprocess, where a biomass generating growth stage is followed by a production stage, where key enzyme levels are dynamically reduced. The limitation of a macronutrient can be used to “switch” cellular metabolism from growth to production. (b) strain variants are screened in high throughput screens. The best producers identified from screening are predictably and rapidly scaled to (d) larger instrumented bioreactors, and (e) subsequently to industrially relevant levels. Importantly this workflow is greatly enhanced by reducing the need for significant process optimization and resulting iteration (red arrows). Reduction in cycle numbers and the need for process optimization is a result of process standardization (a) as well as increased strain robustness and scalability (f-i). (f) During growth the metabolic network in the cell is regulated by internal and environmental variables. (Black circles represent metabolites and black lines represent enzymatic reactions.) In many cases regulation is mediated through metabolic crosstalk in central metabolism where key metabolites control other enzymes sometimes through feedback inhibition (red lines) (g) Upon dynamic control levels of central enzymes reduced leading to reduction in regulator metabolites reducing feedback regulation of central metabolism. This not only enables improved flux through central metabolism and product formation (green circles and lines represent a heterologous production pathway induced in stationary phase) but also (h) robustness or insensitivity to environmental variables. Process robustness leads to (i and b-d) improved process and strain scalability.

Additionally, in these studies, the initial scale up of the best producing strains from microfermetnation based screens to standardized two-stage processes in instrumented bioreactors was successful and did not require significant process optimization. As depicted in Figure 1, one of our central hypotheses to explain this rapid scale up was that by deregulating metabolism in stationary phase, strain performance was more robust to environmental (process) conditions. Simply put, the dynamically deregulated metabolic network has a more limited ability to respond, or less sensitivity to the environment. In this study we test this hypothesis using strains and bioprocess for alanine synthesis. L-alanine, one of the simplest amino acids, has a variety of uses ranging from those in human nutrition, and as a precursor to several pharmaceuticals, to larger volume applications such as an intermediate in the production of the biodegradable, environmentally friendly chelator, methylglycinediacetic acid (MGDA).^32–34^ Biochemically, L-alanine can be produced from pyruvate via a single biochemical step and has previously been produced via fermentation both aerobically and anaerobically in a variety of hosts, including *E. coli.^35–37^* Previous aerobic production of L-alanine has required significant process optimization, even in two-stage approaches, making this amino acid an excellent test case to evaluate the impact of two-stage dynamic control on robustness and scalability. ^35–37^

## Results

### Strain Design

To produce alanine we leveraged a previously reported L-alanine dehydrogenase from *B. subtili*s with mutations altering the cofactor specificity (to prefer NADPH (*ald*\*)). ^38^ As illustrated in Figure 2a, this reversible enzyme converts 1 mole each of pyruvate, ammonia and NADPH into 1 mole of alanine. The *ald*\* was expressed on a high copy plasmid using the robust low phosphate inducible *yibD* gene promoter (yibDp).^17^ Additionally, in several strains the native *E. coli* alanine exporter (*alaE*) was also expressed. Refer to Supplemental Tables S1 and S2 for a list of plasmids and strains used in this study.

**Figure 2:**
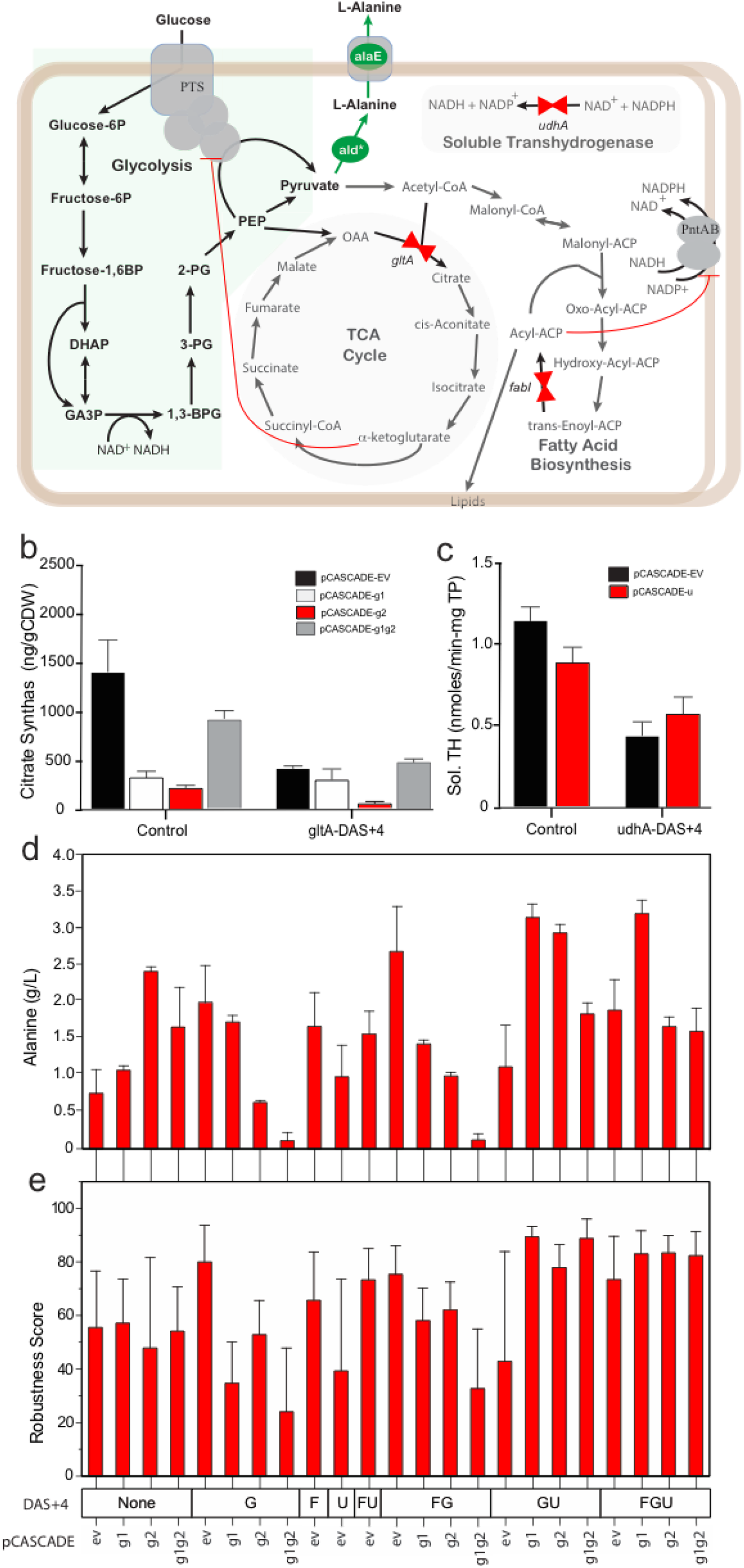
Metabolic pathways engineered for alanine production using two-stage dynamic control. (a) Central Metabolism: Glycolysis, the Citric Acid Cycle (TCA), and Fatty Acid Biosynthesis, and the Soluble Transhydrogenase. Key valve candidate enzymes/genes that are “turned off” to reduce flux through central metabolism are highlighted with red arrows and include: citrate synthase (*gltA-“G”*), enoyl-ACP reductase (*fabI-“F”*), and the soluble transhydrogenase (*udhA-“U”*). Enzymes for that are dynamically “turned on” are highlighted by green arrows and include an NADPH-dependent alanine dehydrogenase (*ald*\*), and alanine exporter (*alaE*). Importantly, dynamic reduction in citrate synthase levels reduce alpha-ketoglutarate pools which inhibit PTS dependent glucose uptake,^22^ and reduced fatty acid metabolites alleviate inhibition of the membrane bound transhydrogenase (PntAB). ^21^ (b-c) Dynamic control over citrate synthase (b) and soluble transhydrogenase (c) levels in two-stage microfermentations. (b) Citrate synthase levels due to proteolysis and/or gene silencing were measured via an ELISA measuring the levels of a C-terminal GFP-tag. Data for the ev and g2 silencing gRNAs was taken from Li et al. ^22^ (c) Changes in soluble transhydrogenase levels due to dynamic control were measured using an enzymatic assay on crude lysates. (mg TP = mg of total lysate protein). (d-e) Results of two-stage microfermentations producing alanine. (d) Average alanine titer and (e) robustness (RS3) in two-stage production with strains having different proteolysis and silencing combinations, n=3. Abbreviation: PTS –glucose phosphotransferase transport system, P – phosphate, BP-bisphosphate, OAA – oxaloacetate, DHAP-dihydroxyacetone phosphate, GA3P-glyceraldehyde-3-phosphate, 1,3-BPG – 1,3 bisphosphoglycerate, 3-PG – 3-phosphoglycerate, 2-PG – 2-phosphoglycerate, PEP-phosphoenolpyruvate, MSA – malonate semialdehyde, ACP – acyl carrier protein,

In addition to the pathway to produce alanine, strains were also modified with a series of metabolic valves. As mentioned above we have previously reported the identification of valves which lead to deregulation of glucose uptake (GltA), acetyl-CoA flux (Zwf), and NADPH synthesis through the membrane bound transhydrogenase (FabI). ^21,22^ Additionally, we have reported that proteolytic degradation of the soluble transhydrogenase (UdhA) can improve NADPH pools. ^21^ As depicted in Figure 2a, alanine production requires pyruvate and NADPH, so we designed strains with combinations of valves reducing levels of GltA (“G” Valves), FabI (“F” Valves), and UdhA (“G” Valves). Firstly, we evaluated the impact of silencing both promoters driving *gltA* expression. While we previously have reported the impact of GltA proteolysis and silencing of the gltAp2 promoter (pCASCADE-g2, which is closer to the transcription start site) on citrate synthase levels, here we additionally evaluated the impact of silencing the gltAp1 promoter (pCASCADE-g1). Additionally, we confirmed the impact of proteolysis and silencing of the soluble transhydrogenase, UdhA, (previous results were performed using xylose, rather than glucose based media ^21^). Results are given in Figure 2b and c. As previously reported, proteolytic degradation of UdhA alone had the largest impact on soluble transhydrogenase levels and will constitute the “U valve” in the remainder of this study. In the case of GltA, we were were surprised that while proteolysis of GltA as well as silencing of either the gltAp1 or gltAp2 promoter led to decreases in citrate synthase levels, the combination of gRNAs to silence both promoters actually resulted in an increase in activity levels (Figure 2d). More work is needed to understand this phenomenon.

### Alanine Production in Micro-Fermentations

This panel of alanine “Valve” strains were constructed and evaluated for alanine production in standardized, two-stage, 96-well plate based, minimal media micro-fermentations as reported by Moreb et al. ^17^ Alanine titers after 24 hours of production are given in Figure 2d. Briefly, titers ranged from ~ 0.2 g/L to ~ 3.5 g/L of alanine (per 1 unit of optical density, OD_600nm_) and varied significantly with respect to the valves. As Figure 2d demonstrates, several valve combinations led to improved performance when compared to the DLF_Z0025 control with no valves. Generally the best performing strains had “F”, “G” and “U” valves, which is consistent with the previously reported effects of these valves. ^21,22^ Aditional studies are required to better understand how the variations in multiple enzyme activities contribute to production.

### Micro-fermentation Robustness

In order to assess robustness, these “Valve” strains were evaluated under several different micro-fermentation process conditions. Glucose concentration and oxygen delivery (key process variables impacting strain performance in traditional aerobic fermentations ^11^) were varied. We then developed scores to quantify environmental/process robustness (Equations (1), (2), (3) and (4)). In these equations, *RS0*_*ij*_ refers to the robustness of strain *i* under process condition *j*. *RS0*_*ij*_ is effectively 1 minus the relative deviation. Subtraction was chosen to give a robustness score of 100, if no error is observed. *RS1*_*i*_ is the average robustness for strain *i* (average *RS0*_*ij*_, where m is the total number of process conditions) under all process conditions. While the average relative standard error (RS1) may seem like a good candidate to assess robustness, this metric reflects the average and does not indicate the largest deviation from the mean performance. We also wanted to capture the largest deviation, so we developed the robustness score *RS2. RS2*_*i*_ is the minimum robustness (minimum *RS0*_*ij*_) for strain *i* under all process conditions. RS2 is a more strict measure of robustness and measures the maximal deviation a strain has under any given process condition. Our final estimate of robustness is RS3, the average of the mean and minimal robustness scores. Larger RS1, RS2 and RS3 scores all indicate more robust strains. An RS1, RS2 or RS3=100 would have an identical alanine titer under all conditions evaluated and be perfectly robust. More variability in titer across conditions would result in lower scores.

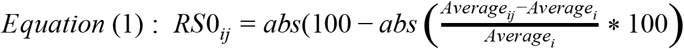

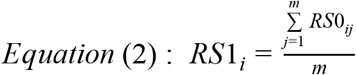

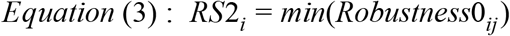

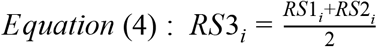

The robustness (RS3) of all valve strains are given in Figure 2e. (Refer to Supplemental Table S6 for details on process conditions. RS1 and RS2 scores are given in Supplemental Table S7). Next, to compare environmental robustness of the two-stage approach enabled by dynamic control to a more traditional growth associated processes, we also constructed five strains, with constitutively expressed alanine dehydrogenase (*ald*\*), capable of the growth associated (as well as stationary phase) production of alanine. These growth associated strains varied in the strength of the promoter used to drive *ald*\* expression,^39^ yet utilized the same common non-valve host (DLF_Z0025, Supplemental Materials Table S5). Results comparing two-stage DMC with growth associated production are given in Figure 3.

**Figure 3:**
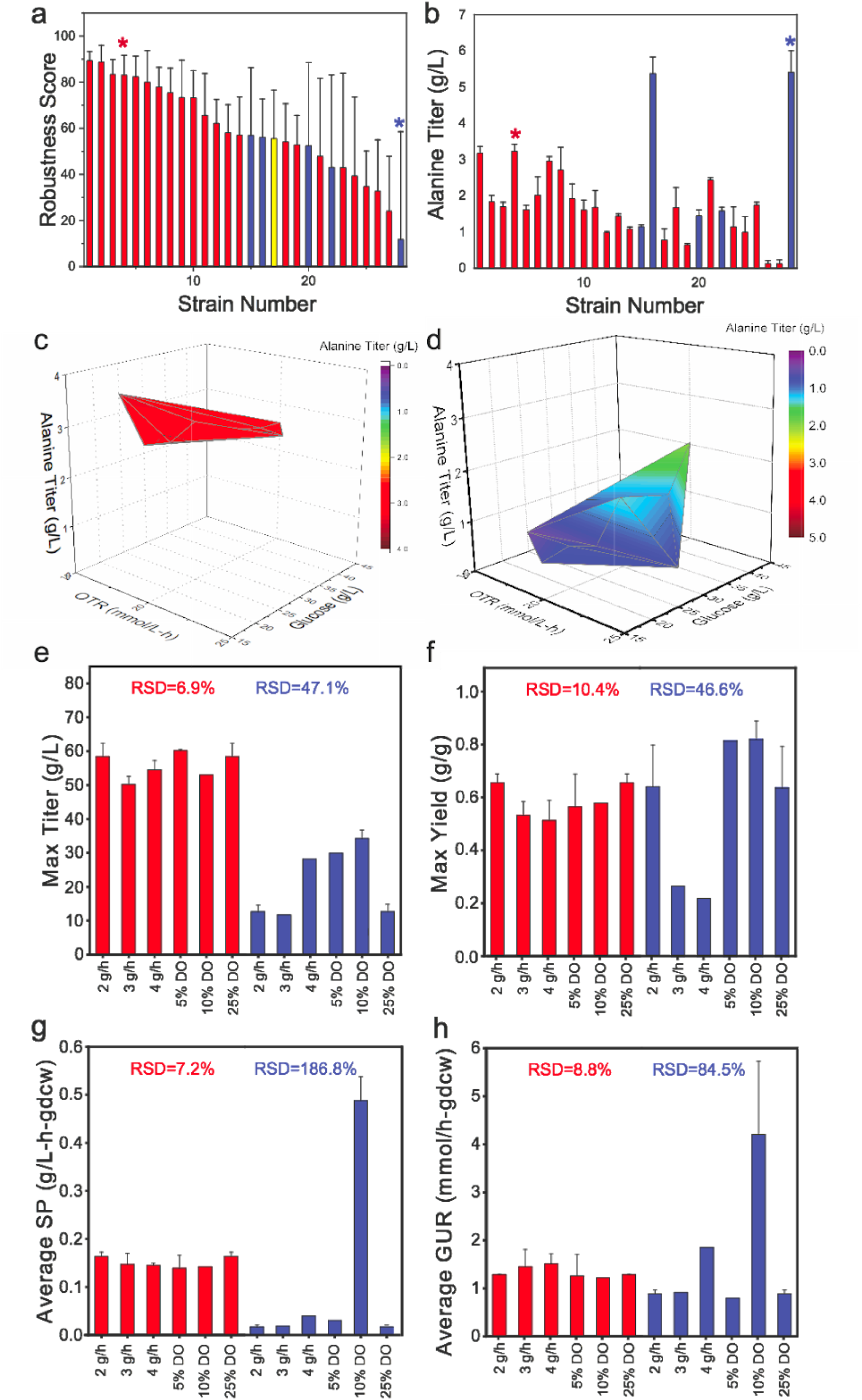
Robustness Comparison between two stage and growth associated approaches. (a) Rank order of the RS3 scores for all alanine strains evaluated, red bars indicate valve alanine strains, and blue bars indicate growth associated (GA) alanine strains, yellow bar is the control strain DLF_Z0025. (b) Alanine titer for all strains (g//L) in the same rank order as panel (a). For alanine titers in panels (a) and (b) the asterisks indicate strains chosen for subsequent robustness studies. (c and d) Microfermentation titer of (c) the dynamic control strain (Silencing: g1, Proteolysis: “FGU”), and (d) strain GA2 as a function of batch glucose and OTR. (e through h) A comparison of process robustness between two-stage (red) and growth associated (blue) processes in instrumented 1L bioreactors, using the “Valve” alanine (Silencing: g1, Proteolysis: “FGU”) strain, and growth associated strain GA2. (e) maximum titer, (f) maximum yield, (g) average specific productivity (SP) and (h) average glucose uptake rate (GUR) are compared for the two approaches in different processes with variations in feed rate (glucose) and dissolved oxygen (D.O.) during production, n=2. Total relative standard deviation (RSD) for all process conditions is given.

The two-stage process (production in stationary phase alone), with no metabolic valves, has a robustness score of ~ 55% (Figure 3a), which was comparable with the most robust growth associated strains. Dynamic control improved robustness in a valve specific manner. The combination of “F”, “G” and “U” valves has consistently improved robustness, which can be explained by the functions of these valves. As mentioned above the “G” valve alleviates inhibition of sugar uptake leading to pyruvate synthesis, and the “F” and “U” valves alleviate inhibition of NADPH production and NADPH consumption, respectively. Pyruvate and NADPH are the two substrates required for alanine synthesis and the elimination of key metabolic regulatory mechanisms controlling their production would be anticipated to increase robustness. Additionally, there was no correlation between robustness and titer for the growth associated strains (Figure 3b-4c).

High throughput approaches are usually used to identify candidate strains that should be moved forward, for example, to evaluation in bioreactors, and this would traditionally be based on a rank order evaluation of titer, which is presented in Figure 3b. In this analysis, the growth associated strain with the highest performance would likely be chosen, without consideration of robustness (Figure 3a). In the case of the growth associated strains tested, the strain with the highest titer also had the lowest robustness score. Improved robustness for “Valve” strains greatly simplifies strain comparisons. Refer to Supplemental Figure S1, and Table S7 for a summary of the robustness data for all strains tested.

### Robustness in Instrumented Fermentations

Robustness of the two-stage DMC and traditional growth associated approach was also assessed at the 1L scale, in fully instrumented bioreactors. Fermentations were performed as described by Menacho-Melgar et al, ^29^ targeting biomass levels of 10 gCDW/L, and varying set point dissolved oxygen levels and glucose feed rates during the production stage (post phosphate depletion). A single robust “Valve” strain (with proteolysis of the *fabI* (“F”), *gltA* (“G”) and *udhA* (“U”) gene products and silencing of the gltAp1 promoter (g1)) and the growth associated strain with the highest micro-fermentation titer (Strain “GA2” from Figure 3b) were compared. The response of these strains to process conditions in microfermentations are given in Figure 3c and d respectively. In fermentations, the “Valve” strains showed consistent performance in all process conditions evaluated (Figure 3e-h), with a maximum relative standard deviation (RSD) of less than 11%, consistent with results from micro-fermentations. Comparatively, the maximum RSD for the growth associated strain was close to 200%. This growth associated strain showed reduced performance under low OTR and high glucose concentrations in micro-fermentations, while at the 1L scale, lower feed rates (lower residual glucose concentrations) and higher dissolved oxygen concentrations led to reduced performance. Although the best performing growth associated strain had a higher alanine titer than any “Valve” strain in micro-fermentations (Figure 3b), the maximal titer for the “Valve” strain was almost two-fold higher than the growth associated strain when evaluated at the 1L scale (Figure 3e).

### Initial Strain Commercialization & Pilot Scale Production

With these initial results including both reasonable titers and robustness, we next turned to the initial commercialization of this alanine strain/process, and performed an initial intellectual property landscape assessment. We found one issued patent with claims covering strains with the combined deletion/disruption of four genes: *ldhA*, *adhE* and *ack-pta*.^*40*^ Although the claims broadly cover any strain having these four gene modifications, the original technology was aimed at optimizing anaerobic succinate production in *E. coli*. ^41^ While these four modifications are in our starting host strain (DLF_Z0025), our process is aerobic rather than anaerobic and we sought to see if a deletion of *adhE* (which is responsible for ethanol production under anaerobic conditions ^42^) was necessary for optimal aerobic stationary phase alanine synthesis in our “FGU” valve background. Toward this end we constructed a derivative of the “FGU” host where the *adhE* gene was reintroduced (+adhE) and measured alanine in microfermentations compared to its parent. Results are given in Figure 4a. To our surprise, the reintroduction of the *adhE* gene actually led to an increase in alanine production. The mechanism underlying this is unclear, but one hypothesis may be the ability of AdhE to scavenge any low levels of toxic acetaldehyde which may be produced into acetyl-CoA,^43^ although further work is needed to investigate this result. We then reevaluated the impact of additional silencing of *gltA* expression in the “FGU”+adhE background, with overexpression of *ald*\* and the alanine exporter *alaE*, and in this host silencing of the gltAp2 promoter (g2) led to the highest alanine titers (Figure 4b). This strain (“FGU”+adhE, pCASCADE-g2, pSMART-Ala10) was then scaled to instrumented bioreactors (Figure 4c) and then to pilot scale production (Figure 4d). Lastly, we turned to intensify the process by increasing biomass levels to ~ 20gCDW/L resulting in alanine titers of 100-120g/L in effectively a 45 hr fermentation, (accounting for the productive period that would be used at larger scales with optimized seed trains). The overall yield in this process was 0.74 g alanine /g of glucose, with the yield in the production phase reaching 0.81 g alanine / g of glucose.

**Figure 4:**
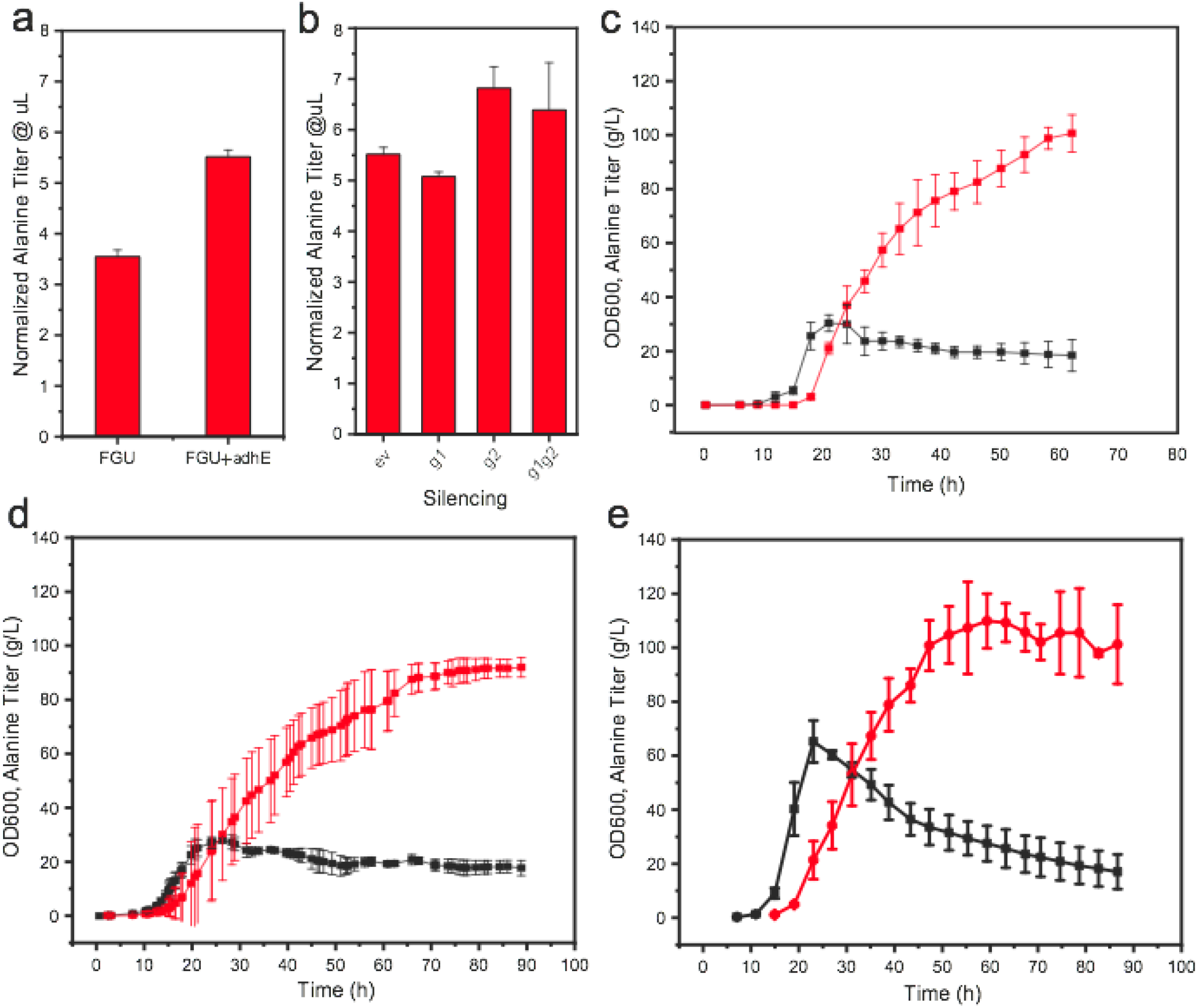
Initial strain commercialization and scale up. All strains contain the high copy plasmid with the low phosphate inducible expression of *ald*\* and *alaE* (pSMART-ala10). (a) Comparison of the micro-fermentation alanine titer for the “FGU” valve host (which contains an *adhE* gene deletion) with the “FGU” valve host where the *adhE* gene was reintroduced (FGU+adhE). (b) Alanine titers in microfermentations using the FGU+adhE host with a set of pCASCADE plasmids silencing *gltA* gene expression (ev - empty vector). (c-e) Biomass (black) and alanine (red) production in fermentations using host strain FGU+adhE with pCASCADE-g2 in (c) 6L lab scale bioreactors, (d) a pilot scale 4000L bioreactor and (e) 6L lab scale bioreactors, with increased biomass levels.

### Scalability of Microfermentations

With the predictable scale up from lab to pilot scale reactors, we next assessed the impact of increased environmental robustness on scalability from the *μ*L scale to L scale fermentations. To evaluate the scalability of the system, “Valve” alanine strains with statistically differentiated performance in micro-fermentations (P-value < 0.001) were evaluated in standardized two-stage 1 L fermentations (Supplemental Materials, Table S8) A standardized phosphate limited two-stage fermentation protocol, targeting biomass levels of 10gCDW/L, and controlled at a 25% dissolved oxygen setpoint was utilized for strain evaluation. This protocol yields highly reproducible growth stage results, with minimal strain to strain variability. More significant variability was observed during the production stage as a result of differing base utilization by different strains. This consistency is contrasted to the more variable growth of growth associated production strains (Supplemental Materials Figure S2). To date, statistically different performances observed in micro-fermentations have scaled predictably to 1L fermentations. This is a stark contrast to the growth associated strains. Data are summarized in Figure 5. Having demonstrated that the two-stage approach had improved scalability in the case of alanine, we assessed the scalability of strains we engineered for the two-stage production of citramalate and xylitol, several of which we have recently reported ^21,22^ In the case of citramalate, the combinations of reductions in citrate synthase (GltA) levels and glucose-6-phosphate dehydrogenase (Zwf) levels led to deregulation of an improved acetyl-CoA flux through pyruvate ferredoxin oxidoreductase and improved citramalate synthesis. ^22^ In the case of xylitol, the combination of decreases in Zwf levels along with reduced FabI levels led to improved NADPH flux both through pntAB and via pyruvate ferredoxin oxidoreductase coupled with NADPH dependent ferredoxin reductase acting as a dehydrogenase. ^21^ Scalability, results are given in Figure 6. In both cases, again improved performance in microfermentations correlated with improved performance in instrumented bioreactors.

**Figure 5:**
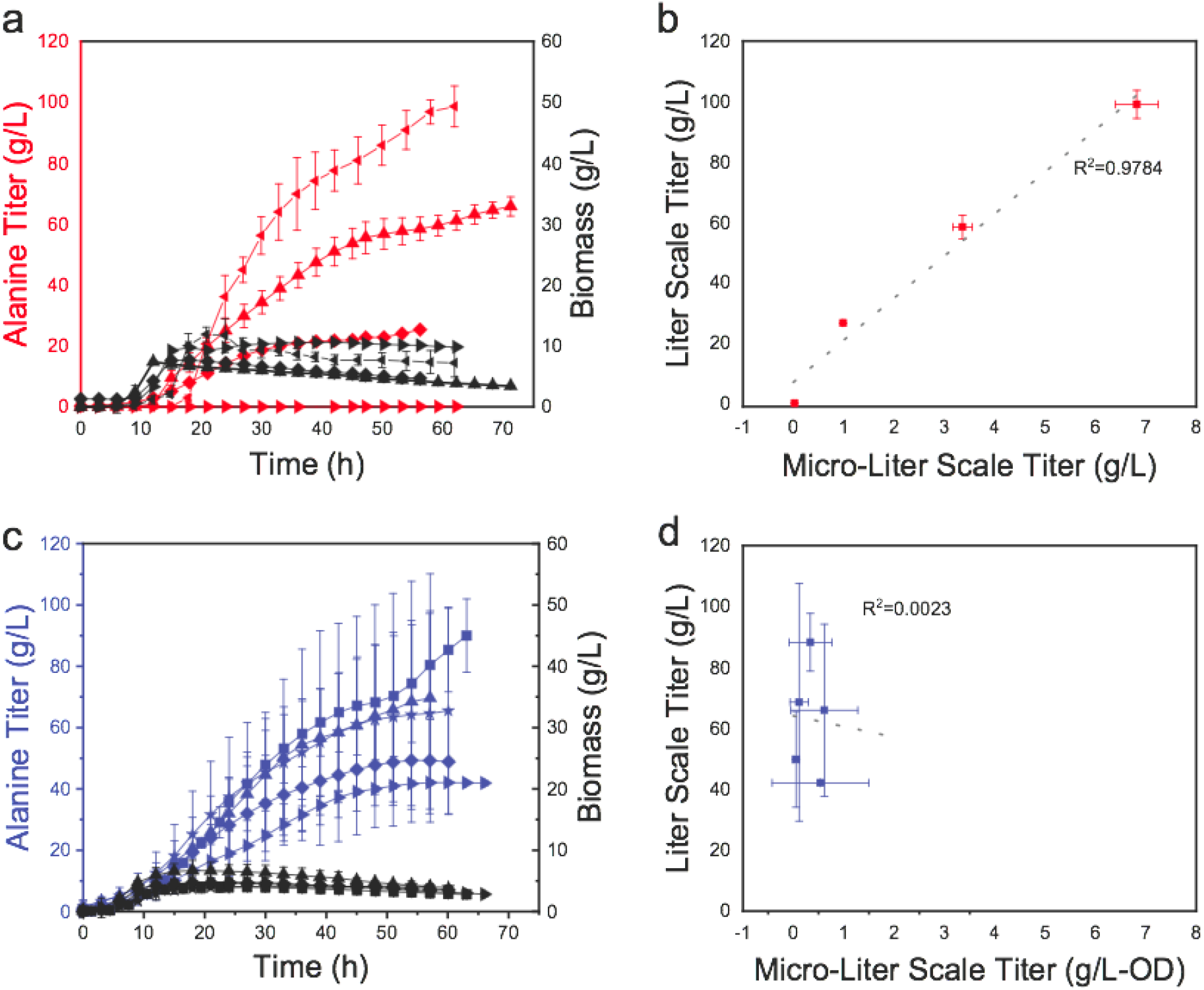
μL to 1L scalability plot for two-stage “Valve” and growth associated alanine producing strains. Strains used for this plot are listed in Supplemental Materials, Table S8. A comparison of the performance of (a) “Valve” and (c) growth associated strains in standard fermentation conditions (2 g/h feed rates and 25% D.O.) at 1L scale. Black squares, OD600 for strain, red or blue squares are alanine titers. (b&d) Scalability, or comparison of the specific alanine production in microfermentations to that achieved in instrumented reactors. (b) two-stage “Valve” strains and (d) growth associated strains. All microfermetnation (μL) data are from n >=3 replicates and fermentation data are n>=2 replicates.

**Figure 6:**
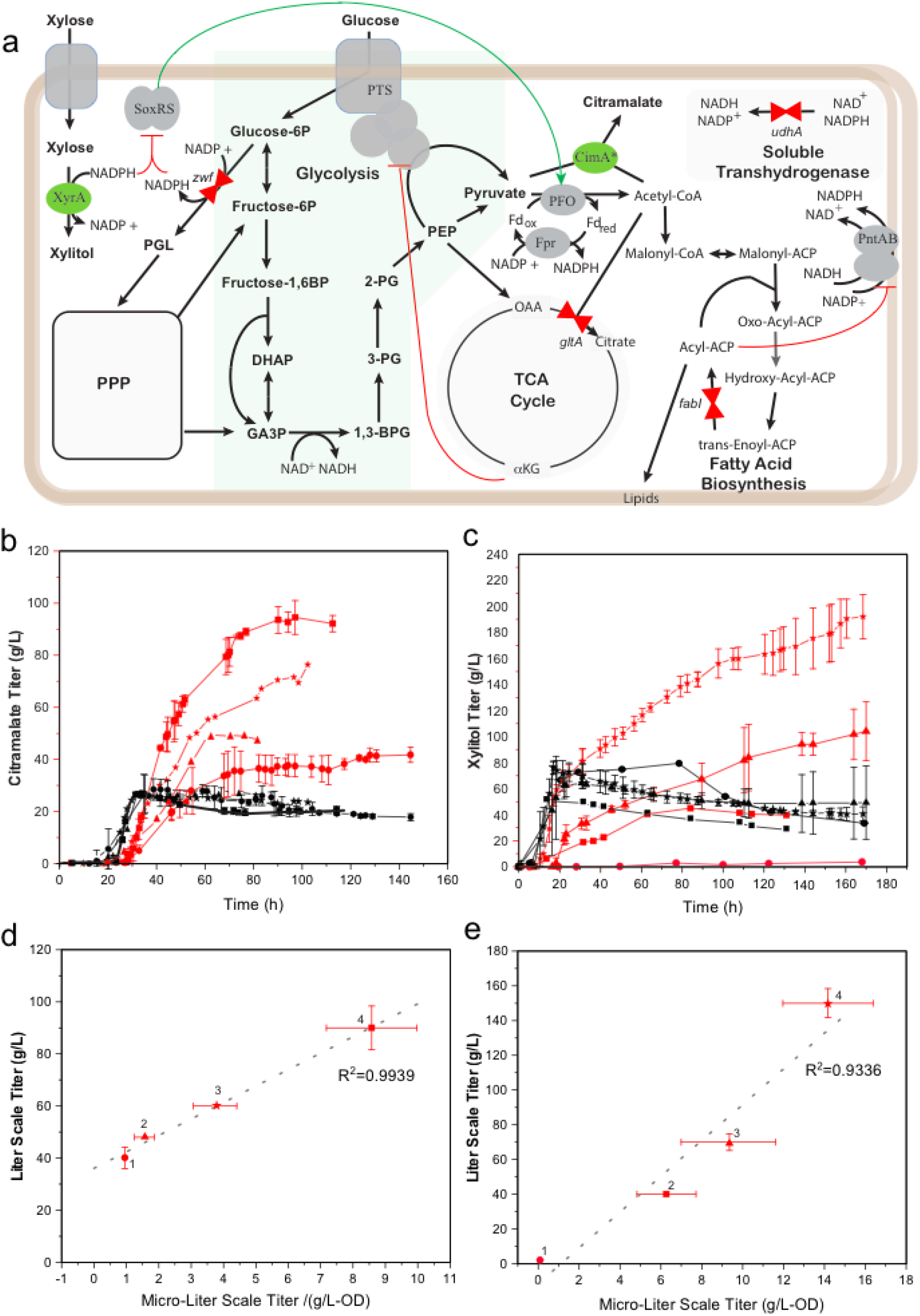
Scalability of citramalate and xylitol producing strains leveraging two-stage dynamic control. (a) An overview of the pertinent metabolism for citramalate and xylitol production. Citramalate is produced by a single enzyme, a mutant citramalate synthase (CimA* encoded by the *cimA3.7* gene ^22^) from one mole of pyruvate and one mole of acetyl-CoA. Xylitol is produced by a single enzyme xylose reductase (XyrA^21^). Valves are as described in Figure 2, with the addition of a metabolic valve (both proteolysis and gene silencing) reducing levels of glucose-6-phosphate dehydrogenase (Zwf). Two-stage fermentation profiles (black lines OD600nm and red lines product titers) for (b) citramalate producing strains and (c) xylitol producing strains. (d-e) Plots of microfermentaton performance vs. fermentation performance for (d) citramalate and (e) xylitol producing strains. Fermentation titers are taken as the maximal titer within 60 hrs after the onset of stationary phase. Citramalate producing strain information is as follows: Strain 1: Proteolysis none, pCASCADE-ev, pHCKan-yibDp-cimA3.7 Strain 2: Proteolysis F,G,U, pCASCADE-g1g2z, pHCKan-yibDp-cimA3.7, Strain 3: Proteolysis G, pCASCADE-g2, pHCKan-yibDp-cimA3.7, Strain 4: Proteolysis GZ, pCASCADE-g2z, pHCKan-yibDp-cimA3.7. Xylitol producing strain information is as follows: Strain 1: Proteolysis none, pCASCADE-ev, pHCKan-xyrA, Strain 2: Proteolysis Z, pCASCADE-g1z, pHCKan-xyrA, Strain 3: Proteolysis FZ, pCASCADE-z, pHCKan-xyrA, Strain 4: Proteolysis FZ, pCASCADE-z, pHCKan-xyrA, pCDF-pntAB (which enables additional overexpression of pntAB). Data for citramalate strains 1,3, and 4 was taken from Li et al. ^22^. Data for xylitol strains 1,3, and 4 was taken from Li et al. ^21^). Abbreviation: PPP - pentose phosphate pathway, PTS –glucose phosphotransferase transport system, P – phosphate, BP-bisphosphate, OAA – oxaloacetate, DHAP-dihydroxyacetone phosphate, GA3P-glyceraldehyde-3-phosphate, 1,3-BPG – 1,3 bisphosphoglycerate, 3-PG – 3-phosphoglycerate, 2-PG – 2-phosphoglycerate, PEP-phosphoenolpyruvate, MSA – malonate semialdehyde, ACP – acyl carrier protein,

## Discussion

There is significant precedent in the biotechnology industry for using and scaling of two-stage processes similar to that described, ^8^ including many used for heterologous expression of proteins.^8,44^ This approach is not compatible with growth associated processes, and many more traditional metabolic engineering approaches focused on coupling growth and production. Historically many of the successful metabolic engineering efforts in the production of small molecules have leveraged the power of coupling product formation with growth. This has allowed for the classical design and selection of industrial strains to produce numerous products including ethanol, succinic acid, lactate and isobutanol, which have leveraged the power of evolution and selection, to reach optimal stoichiometric metabolic fluxes in engineered networks. ^45,46^ However, the requirement for growth coupling can also provide serious process constraints and the need for significant process optimization, Two-stage control enables deregulation of metabolism which can both improve process robustness and allow for metabolic states not compatible with cellular growth.

Some readers may be of the opinion that with growth associated production, traditional process optimization is likely to yield improved results, and that we have not pushed the growth associated strains described above to their potential. For example, the optimal performance of any one of the growth associated strains may well exceed the performance measured in these initial experiments. This is indeed true, optimization of various process parameters, feed limitation and oxygen control could all be used and perhaps the optimal performance would exceed any of our current results. Several groups have produced alanine to titers from 80-110g/L in several microbes, even in the context of two-stage production, however in these studies significant process optimization was needed to achieve these results.^35–37^ Similarly previous studies producing citramalate in *E. coli* have required significant process optimization, including the optimization of media, complex feeding strategies and oxygen control. ^47,48^

However this is the process optimization cycle that improved strain robustness can minimize. These optimization cycles can be very time consuming, expensive and strain dependent (leading to unique process solutions), and almost always require experiments using instrumented bioreactors. By engineering robustness or insensitivity to process conditions, a standardized process, like the one described, can be developed independent of product. This enables scalable highthroughut approaches and the evaluation of larger numbers of genetic variants, enabling challenges to be overcome with metabolic engineering strategies rather than process changes. This is critical to remove or minimize the process development bottleneck in bioprocess development and take advantage of the ever increasing speed of synthetic biology. In this framework, robustness as a metric is as important as titer, when considering the need for scale up, either from high throughput studies to bioreactors or from lab scale reactors to pilot scale. In this work we demonstrate that the dynamic two-stage deregulation, one approach to enhance robustness to environmental variables, enables improved scalability of high throughput small-scale studies to larger instrumented fermentations. More generally approaches that improve robustness and make biology more predictable are an important area for future investigation, with the potential to simplify high-throughput experimentation and scale up of novel bioprocesses.

## Methods

### Reagents and Media

Materials and reagents, unless otherwise specified, were purchased from Sigma (St. Louis, MO). C13 labeled Alanine (2,3-13C2, 99%) (Item # CLM-2734-PK) was purchased from Cambridge Isotope Laboratories, Inc. (Tewksbury, MA). Luria Broth (lennox formulation) was used for routine strain and plasmid propagation and construction. Working antibiotic concentrations were as follows: kanamycin (35 μg/mL), chloramphenicol (35 μg/mL), zeocin (50 μg/mL), gentamicin (10 μg/mL), blasticidin (100 μg/mL), puromycin (150 μg/mL). Puromycin selection was performed with LB supplemented with 50 mM potassium phosphate buffer (pH=8.0) to maintain pH for adequate selection. Media SM10++, SM10+, FGM3 No phosphate, SM10 no phosphate, FGM10 and FGM25 were all prepared as previously reported. ^17,29^

### Plasmids and strains

All host strains were constructed as previously reported. (Refer to Supplemental Table S2).^21,22^ Primers used for the design and construction of CASCADE guides arrays were listed in Supplemental Table S3. The pCASCADE guide plasmids pCASCADE-g1 and pCASCADE-g1g2 were prepared by sequentially amplifying complementary halves of each smaller guide plasmid by PCR, followed by subsequent DNA assembly as previously described. ^22^ Alanine production plasmids were constructed using codon optimized (Codon Optimization Tool from the IDT) synthetic DNA and the pSMART-HCKan vector (Lucigen, Middleton, WI). Plasmids were assembled using NEBuilder^®^ HiFi DNA Assembly Master Mix following manufacturer’s protocol (NEB, MA). All plasmid sequences were confirmed by DNA sequencing (Eton Bioscience, NC) and deposited with Addgene. To add adhE back to DLF_Z0047, a purR resistance cassette was first integrated 3’ of *adhE* in *E.coli* BW25113 using standard recombineering methods, adhE-purR was then PCR amplified and transferred to DLF_Z0047 again using standard recombineering protocols.^49^ The recombineering plasmid pSIM5 was a kind gift from Donald Court (NCI, https://redrecombineering.ncifcrf.gov/court-lab.html).^50^ adhE-purR in DMC_HS_01718 was confirmed by PCR and DNA sequencing. Primers and gblock used are listed in Table S4.

### Fermentations

Minimal media microfermentations were performed as previously reported by Moreb et al. ^17^ In the robustness studies variations in batch glucose and or aeration were performed as described in Supplemental Materials (Tables S6). OTR was estimated as a function of shaking speed and fill volume according to Duetz and Witholt.^51^ For growth associated alanine micro-fermentations, during production FGM3 with 40 mM phosphate was used instead of No phosphate FGM3 media.

Fermentations in instrumented bioreactors performed as reported by Menacho-Melgar et al. ^29^ with slight modifications to the feeding profiles to ensure adequate glucose and in the pilot scale minimal residual glucose at the end of the fermentation. For details on feeding profiles refer to Supplemental Materials, Table S9 and S10. Pilot scale fermentations were performed identically to those at lab scale, except that frozen seeds were not used to directly inoculate the pilot reactor. Instead frozen seed vials (as reported by Menacho-Melgar et al. ^29^), were used to inoculate 10L lab bioreactors (with FGM10 media), cells were harvested in mid exponential phase and used immediately to inoculate the larger vessel. In the case of growth associated fermentation processes, FGM10 medium with 40 mM phosphate was used instead of FGM10 medium, which normally has 5mM phosphate.

### Enzyme Quantification

Quantification of citrate synthase levels was performed using clones expressing C-terminally GFP tagged GltA and an ELISA to quantify GFP as described by Li et al.^22^ Soluble transhydrogenase was quantified by enzymatic assays on whole cell lysates as described by Li et al. ^21^

### Glucose Quantification

A UPLC-RI method was developed for the quantification of glucose concentration, using an Acquity H-Class UPLC integrated with a Waters 2414 Refractive Index (RI) detector (Waters Corp., Milford, MA. USA). Chromatographic separation was performed using a Bio-Rad Fast Acid Analysis HPLC Column (100 x 7.8 mm, 9 μm particle size; CAT#: #1250100, Bio-Rad Laboratories, Inc., Hercules, CA) at 65 ℃. 5 mM sulfuric acid was used as the eluent. The isocratic elution was as follows: 0–0.1 min, flow rate increased from 0.4 mL/min to 0.42 mL/min, 0.1–12 min flow rate at 0.48 mL/min. Sample injection volume was 10 μL. UPLC method development was carried out using standard aqueous stock solutions of analyte. Peak integration and further analysis was performed using MassLynx v4.1 software. The linear range used for glucose was 1-10 g/L. Samples were diluted as needed to be within the accurate linear range. Dilution was performed using ultrapure water.

### Product Quantification

Two orthogonal methods were used for alanine quantification. First, a reverse phase UPLC-MS/MS method was developed for alanine. Chromatographic separation was performed using a Restek Ultra AQ C18 column (150 mm × 2.1 i.d., 3 μm; CAT#: 9178362, Restek Corporation, Bellefonte, PA) at 70 ℃. The following eluents were used: solvent A: H_2_O, 0.2% formic acid and 0.05% ammonium (v/v); solvent B: MeOH, 0.2% formic acid and 0.05% ammonium (v/v). The gradient elution was as follows: 0–0.1 min isocratic 5% B, flow rate increased from 0.65 mL/min to 0.75 mL/min; 0.1–0.3 min, linear from 5% to 95% B at 0.75 mL/min; 0.3–0.9 min isocratic 95% B at 0.75 mL/min; and 0.9–1.2 min linear from 95% to 5% B at 0.75 mL/min; 1.2–1.3 min isocratic 5% B at 0.75 mL/min. Sample injection volume was 5 μL. UPLC method development was carried out using standard aqueous stock solutions of analyte. Separations were performed using an Acquity H-Class UPLC integrated with a Xevo™ TQD Mass spectrometer (Waters Corp., Milford, MA. USA). Alanine (2,3-13C2, 99%) was used as internal standard for alanine at a concentration of 5 mg/L. MS/MS parameters including MRM transitions were as follows: Alanine 89.95>44.08, CV 15, CE 9; C13-Alanine 91.95>46.08, CV 15, CE 9. Peak integration and further analysis was performed using MassLynx v4.1 software. The linear range for alanine was 1-100 mg/L. Samples were diluted as needed to be within the accurate linear range. Dilution was performed using ultrapure water, and the final 10-fold dilution was performed using solvent A, with 5 mg/L of C13 alanine (2,3-13C2, 99%).

Secondly, an HPLC-UV method was used to quantify L-alanine, Chromatographic separation was performed using a Chirex 3126 (D)-penicillamine column (150 x 4.6 mm, 5 μm; Phenomenex Inc., Torrance, CA) at 50 ℃. 2 mM Copper Sulfate was used as the eluent. An isocratic elution was maintained for 10 minutes per sample at flow rate of 0.75 mL/min. The sample injection volume was 10 μL. Absorbance was monitored at 254 nm. L-alanine eluted at 4.93 minutes. Citramalate and Xylitol were quantified as previously reported. ^21,22^

## Acknowledgements

We would like to acknowledge the following support: NSF EAGER #1445726, DARPA #HR0011-14-C-0075, and ONR YIP #N00014-16-1-2558, as well as support from DMC Biotechnologies, Inc. We would also like to thank the team at the Bioprocess Pilot Facility (Delft, Netherlands https://www.bpf.eu/) for performing the 4000L pilot fermentations.

## Author Contributions

Z. Ye constructed plasmid and strains, performed micro-fermentation and 1L studies, collected analytical data and analyzed all results. J.N. Hennigan performed enzyme assays. Z. Ye, S. Li, J. Lebeau, and J. Wolf performed fermentations. E.A. Moreb assisted with microfermentations. M. Lynch constructed strains, designed experiments and analyzed results. Z. Ye, J.N. Hennigan, E.A. Moreb, S. Li and M.D. Lynch prepared the manuscript.

## Conflicts of Interest

M.D. Lynch, Z. Ye and J. Wolf have equity or options in DMC Biotechnologies, Inc. M.D. Lynch, J.N. Hennigan, and E.A. Moreb have equity in Roke Biotechnologies, LLC.

## References

1. Cameron, D. E., Bashor, C. J. & Collins, J. J. A brief history of synthetic biology. Nat. Rev. Microbiol. 12, 381–390 (2014).

2. Jarboe, L. R. et al. Metabolic engineering for production of biorenewable fuels and chemicals: contributions of synthetic biology. J. Biomed. Biotechnol. 2010, 761042 (2010).

3. Lee, J. W. et al. Systems metabolic engineering of microorganisms for natural and non-natural chemicals. Nat. Chem. Biol. 8, 536–546 (2012).

4. Choi, S. Y. et al. One-step fermentative production of poly(lactate-co-glycolate) from carbohydrates in Escherichia coli. Nat. Biotechnol. 34, 435–440 (2016).

5. Dellomonaco, C., Clomburg, J. M., Miller, E. N. & Gonzalez, R. Engineered reversal of the β-oxidation cycle for the synthesis of fuels and chemicals. Nature 476, 355–359 (2011).

6. Kim, S., Clomburg, J. M. & Gonzalez, R. Synthesis of medium-chain length (C6--C10) fuels and chemicals via β-oxidation reversal in Escherichia coli. J. Ind. Microbiol. Biotechnol. 42, 465–475 (2015).

7. Meadows, A. L. et al. Rewriting yeast central carbon metabolism for industrial isoprenoid production. Nature 537, 694–697 (2016).

8. Burg, J. M., Cooper, C. B., Ye, Z., Reed, B. R. & Moreb, E. A. Large-scale bioprocess competitiveness: the potential of dynamic metabolic control in two-stage fermentations. Current opinion in (2016).

9. Logue, J. B., Findlay, S. E. G. & Comte, J. Editorial: Microbial Responses to Environmental Changes. Frontiers in Microbiology vol. 6 (2015).

10. Waegeman, H. et al. Effect of iclR and arcA knockouts on biomass formation and metabolic fluxes in Escherichia coli K12 and its implications on understanding the metabolism of Escherichia coli BL21 (DE3). BMC Microbiol. 11, 70 (2011).

11. Garcia-Ochoa, F. & Gomez, E. Bioreactor scale-up and oxygen transfer rate in microbial processes: an overview. Biotechnol. Adv. 27, 153–176 (2009).

12. Formenti, L. R. et al. Challenges in industrial fermentation technology research. Biotechnol. J. 9, 727–738 (2014).

13. Shalel Levanon, S., San, K.-Y. & Bennett, G. N. Effect of oxygen on the Escherichia coli ArcA and FNR regulation systems and metabolic responses. Biotechnol. Bioeng. 89, 556–564 (2005).

14. Hemmerich, J. et al. Comprehensive clone screening and evaluation of fed-batch strategies in a microbioreactor and lab scale stirred tank bioreactor system: application on Pichia pastoris producing Rhizopus oryzae lipase. Microbial Cell Factories vol. 13 36 (2014).

15. Ramirez-Vargas, R., Vital-Jacome, M., Camacho-Perez, E., Hubbard, L. & Thalasso, F. Characterization of oxygen transfer in a 24-well microbioreactor system and potential respirometric applications. J. Biotechnol. 186, 58–65 (2014).

16. Huber, R., Roth, S., Rahmen, N. & Buchs, J. Utilizing high-throughput experimentation to enhance specific productivity of an E.coli T7 expression system by phosphate limitation. BMC Biotechnol. 11, 22 (2011).

17. Moreb, E. A. et al. Media Robustness and scalability of phosphate regulated promoters useful for two-stage autoinduction in E. coli. ACS Synthetic Biology (2020) doi:10.1021/acssynbio.0c00182.

18. Lynch, M. D. Into new territory: improved microbial synthesis through engineering of the essential metabolic network. Curr. Opin. Biotechnol. 38, 106–111 (2016).

19. Brockman, I. M. & Prather, K. L. J. Dynamic metabolic engineering: New strategies for developing responsive cell factories. Biotechnol. J. 10, 1360–1369 (2015).

20. Brockman, I. M. & Prather, K. L. J. Dynamic knockdown of E. coli central metabolism for redirecting fluxes of primary metabolites. Metab. Eng. 28, 104–113 (2015).

21. Li, S. et al. Dynamic control over feedback regulatory mechanisms improves NADPH fluxes and xylitol biosynthesis in engineered E. coli. bioRxiv 2020.07.27.222588 (2020) doi:10.1101/2020.07.27.222588.

22. Li, S., Ye, Z., Lebeau, J., Moreb, E. A. & Lynch, M. D. Dynamic control over feedback regulation identifies pyruvate-ferredoxin oxidoreductase as a central metabolic enzyme in stationary phase E. coli. bioRxiv 2020.07.26.219949 (2020) doi:10.1101/2020.07.26.219949.

23. Ye, Z., Lynch,M.D., Trahan, A.D., Rodriguez, D.L., Cooper, C.B. Bozdag, A. Compositions and methods for rapid and dynamic flux control using synthetic metabolic valves. Patent (2015).

24. Venayak, N., von Kamp, A., Klamt, S. & Mahadevan, R. MoVE identifies metabolic valves to switch between phenotypic states. Nat. Commun. 9, 5332 (2018).

25. Soma, Y., Tsuruno, K., Wada, M., Yokota, A. & Hanai, T. Metabolic flux redirection from a central metabolic pathway toward a synthetic pathway using a metabolic toggle switch. Metab. Eng. 23, 175–184 (2014).

26. Sander, T., Wang, C. Y., Glatter, T. & Link, H. CRISPRi-Based Downregulation of Transcriptional Feedback Improves Growth and Metabolism of Arginine Overproducing E. coli. ACS Synth. Biol. 8, 1983–1990 (2019).

27. Venayak, N., Raj, K., Jaydeep, R. & Mahadevan, R. An Optimized Bistable Metabolic Switch To Decouple Phenotypic States during Anaerobic Fermentation. ACS Synth. Biol. 7, 2854–2866 (2018).

28. Lynch, M. D., Gill, R. T. & Lipscomb, T. E. W. Method for producing 3-hydroxypropionic acid and other products. US Patent (2016).

29. Menacho-Melgar, R. et al. Scalable, two-stage, autoinduction of recombinant protein expression in E. coli utilizing phosphate depletion. Biotechnol. Bioeng. 26, 44 (2020).

30. McGinness, K. E., Baker, T. A. & Sauer, R. T. Engineering controllable protein degradation. Mol. Cell 22, 701–707 (2006).

31. Luo, M. L., Mullis, A. S., Leenay, R. T. & Beisel, C. L. Repurposing endogenous type I CRISPR-Cas systems for programmable gene repression. Nucleic Acids Res. 43, 674–681 (2015).

32. Schneider, J. et al. Use of glycine-N,N-diacetic acid derivatives as biodegradable complexing agents for alkaline earth metal ions and heavy metal ions and process for the preparation thereof. US Patent (1998).

33. Greindl, T. et al. Process for the preparation of glycine-N, N-diacetic acid derivatives. US Patent (1998).

34. Williams, D. R. Speciation of Chelating Agents and Principles for Global Environmental Management. Biogeochemistry of Chelating Agents 20–49 (2005) doi:10.1021/bk-2005-0910.ch002.

35. Zhang, X., Jantama, K., Moore, J. C., Shanmugam, K. T. & Ingram, L. O. Production of L-alanine by metabolically engineered Escherichia coli. Appl. Microbiol. Biotechnol. 77, 355–366 (2007).

36. Smith, G. M., Lee, S. A., Reilly, K. C., Eiteman, M. A. & Altman, E. Fed-batch two-phase production of alanine by a metabolically engineered Escherichia coli. Biotechnol. Lett. 28, 1695–1700 (2006).

37. Zhou, L., Deng, C., Cui, W.-J., Liu, Z.-M. & Zhou, Z.-M. Efficient L-Alanine Production by a Thermo-Regulated Switch in Escherichia coli. Appl. Biochem. Biotechnol. 178, 324–337 (2016).

38. Lerchner, A., Jarasch, A. & Skerra, A. Engineering of alanine dehydrogenase from Bacillus subtilis for novel cofactor specificity. Biotechnol. Appl. Biochem. 63, 616–624 (2016).

39. Davis, J. H., Rubin, A. J. & Sauer, R. T. Design, construction and characterization of a set of insulated bacterial promoters. Nucleic Acids Res. 39, 1131–1141 (2011).

40. Ka-Yiu, S., Bennett, G. N. & Sanchez, A. Mutant E. coli strain with increased succinic acid production. US Patent (2007).

41. Sánchez, A. M., Bennett, G. N. & San, K.-Y. Efficient Succinic Acid Production from Glucose through Overexpression of Pyruvate Carboxylase in an Escherichia coli Alcohol Dehydrogenase and Lactate Dehydrogenase Mutant. Biotechnology Progress vol. 21 358–365 (2008).

42. Chen, Y. M. & Lin, E. C. Regulation of the adhE gene, which encodes ethanol dehydrogenase in Escherichia coli. J. Bacteriol. 173, 8009–8013 (1991).

43. Rudolph, F. B., Purich, D. L. & Fromm, H. J. Coenzyme A-linked aldehyde dehydrogenase from Escherichia coli. I. Partial purification, properties, and kinetic studies of the enzyme. J. Biol. Chem. 243, 5539–5545 (1968).

44. Dean, J. & Reddy, P. Metabolic analysis of antibody producing CHO cells in fed-batch production. Biotechnol. Bioeng. 110, 1735–1747 (2013).

45. Orth, J. D., Thiele, I. & Palsson, B. Ø. What is flux balance analysis? Nat. Biotechnol. 28, 245–248 (2010).

46. Yim, H. et al. Metabolic engineering of Escherichia coli for direct production of 1,4-butanediol. Nat. Chem. Biol. 7, 445–452 (2011).

47. Wu, X., Tovilla-Coutiño, D. B. & Eiteman, M. A. Engineered citrate synthase improves citramalic acid generation in Escherichia coli. Biotechnol. Bioeng. 117, 2781–2790 (2020).

48. Webb, J. P. et al. Efficient bio-production of citramalate using an engineered Escherichia coli strain. Microbiology 164, 133–141 (2018).

49. Li, X.-T., Thomason, L. C., Sawitzke, J. A., Costantino, N. & Court, D. L. Positive and negative selection using the tetA-sacB cassette: recombineering and P1 transduction in Escherichia coli. Nucleic Acids Res. 41, e204 (2013).

50. Sharan, S. K., Thomason, L. C., Kuznetsov, S. G. & Court, D. L. Recombineering: a homologous recombination-based method of genetic engineering. Nat. Protoc. 4, 206–223 (2009).

51. Duetz, W. A. & Witholt, B. Oxygen transfer by orbital shaking of square vessels and deepwell microtiter plates of various dimensions. Biochem. Eng. J. 17, 181–185 (2004).

