## Supplementary Materials for "Two-stage Dynamic Deregulation of Metabolism Improves Process Robustness & Scalability in Engineered *E. coli*"

### Supplemental Materials

#### 1. Strains & Plasmids

**Table S1:** List of chromosomally modified strains.

| Strain | Genotype | Source |
|---|---|---|
| BW25113 | F-, λ-, Δ(araD-araB)567, lacZ4787(del)(::rrnB-3) , rph-1, Δ(rhaD-rhaB)568, hsdR514 | CGSC |
| DLF_Z0025 | F-, λ-, Δ(araD-araB)567, lacZ4787(del)(::rrnB-3) , rph-1, Δ(rhaD-rhaB)568, hsdR514, ΔackA-pta, ΔpoxB, ΔpflB, ΔldhA, ΔadhE, ΔiclR, ΔarcA, ΔsspB::frt, Δcas3::tm-ugpb-sspB-pro-casA | [1] |
| DLF_Z0028 | DLF_Z0025, fabI-DAS+4-gentR | [2] |
| DLF_Z0038 | DLF_Z0025, fabI-DAS+4-gentR, udhA-DAS+4-bsdR | [2] |
| DLF_Z0039 | DLF_Z0025, fabI-DAS+4-gentR, gltA-DAS+4-zeoR | [2] |
| DLF_Z0043 | DLF_Z0025, gltA-DAS+4-zeoR | [1] |
| DLF_Z0045 | DLF_Z0025, gltA-DAS+4-zeoR, udhA-DAS+4-bsdR | [2] |
| DLF_Z0047 | DLF_Z0025, fabI-DAS+4-gentR, gltA-DAS+4::zeoR, udhA-DAS+4-bsdR | [2] |
| DLF_Z0763 | DLF_Z0025, udhA-DAS+4-bsdR | [2] |
| DMC_HS_01718 | DLF_Z0047, adhE-purR | This study |

**Table S2:** List of plasmids used in this study.

| **Plasmid Utilized in this Study** | | | |
|---|---|---|---|
| **Plasmid** | **Purpose** | **Source** | **Reference** |
| pSMART-HC-Kan | Backbone Vector | Lucigen | -- |
| **Plasmid Constructed in this Study** | | | |
| **Plasmid** | **Plasmid Name** | **Addgene ID** | **Reference** |
| pSMART-Ala2 | pSMART-HCKan:yibDp-ald* | 71326 | This study |
| pSMART-Ala11 | pSMART-HCKan-GA1-ald* | 87172 | This study |
| pSMART-Ala12 | pSMART-HCKan-GA2-ald* | 87173 | This study |
| pSMART-Ala13 | pSMART-HCKan-GA3-ald* | 87174 | This study |
| pSMART-Ala10 | pSMART-HCKan-yibDp-ald*-alaE | 87135 | [3] |
| pCASCADE-ev | pCASCADE-empty vector control | 65821 | [1] |
| pCASCADE-gltA1 | pCASCADE-gltA1 | 71334 | [1] |
| pCASCADE-gltA2 | pCASCADE-gltA2 | 65817 | [1] |
| pCASCADE-G1G2 | pCASCADE-gltA1-gltA2 | 71348 | This study |

**Table** S3: Details of gRNA construction

| **sgRNA/Primer Name** | **Sequence** | **Template** |
|---|---|---|
| G1G2 | *TCGAGTTCCCCGCGCCAGCGGGGATAAACCG*AAAAGCATATAA<br>TGCGTAAAAGTTATGAAGT*TCGAGTTCCCCGCGCCAGCGGGGA*<br>*TAAACCGT*ATTGACCAATTCATTCGGGACAGTTATTAGT*TCGAG*<br>*TTCCCCGCGCCAGCGGGGATAAACCG* | |
| gltA2-FOR | GCGCCAGCGGGGATAAACCG*TATTGACCAATTCATTC* | pCASCADE-g2 |
| pCASCADE-REV | CTTGCCCGCCTGATGAATGCTCATCCGG | |
| pCASCADE-FOR | CCGGATGAGCATTCATCAGGCGGGCAAG | pCASCADE-g1 |
| gltA1-REV | CGGTTTATCCCCGCTGGCGCGGGGAACTCGAACTTCATAACTT<br>TTAC | |

**Table** S4: Primers and gblock used for DMC_HS_01718 strain.

| **Primer/gblock Name** | **Sequence** |
|---|---|
| AdhE-purR | CGTGAAGCTGGCGTTCAGGAAGCAGACTTCCTGGCGAACGTGGATAAACTGTC<br>TGAAGATGCATTCGATGACCAGTGCACCGGCGCTAACCCGCGTTACCCGCTGAT<br>CTCCGAGCTGAAACAGATTCTGCTGGATACCTACTACGGTCGTGATTATGTAGAA<br>GGTGAAACTGCAGCGAAGAAAGAAGCTGCTCCGGCTAAAGCTGAGAAAAAAG<br>CGAAAAAATCCGCTTAATCCTGACGGATGGCCTTTTTGCGTTTCTACAAACTCTT<br>TTTGTTTATTTTTCTAAATACATTCAAATATGTATCCGCTCATGAGACAATAACCC<br>TGATAAATGCTTCAATAATATTGAAAAAGGAAGAGTATGACTGAATACAAGCCC<br>ACGGTACGCTTGGCGACGCGCGACGATGTTCCCCGCGCTGTTCGTACATTAGCT<br>GCGGCCTTTGCAGATTACCCAGCGACGCGCCATACGGTCGATCCGGACCGCCAT<br>ATCGAGCGTGTCACAGAATTGCAGGAACTTTTCTTAACTCGCGTGGGCCTTGAC<br>ATCGGAAAGGTCTGGGTGGCTGACGATGGCGCTGCAGTGGCTGTTTGGACCAC<br>TCCGGAGAGTGTAGAGGCTGGTGCAGTGTTCGCCGAAATTGGTCCTCGTATGGC<br>CGAATTAAGTGGAAGTCGTCTGGCAGCCCAACAACAAATGGAAGGGTTGCTTG<br>CGCCCCACCGTCCGAAAGAACCCGCGTGGTTCCTTGCCACCGTTGGAGTAAGC<br>CCAGATCACCAGGGGAAGGGTTTAGGATCTGCCGTAGTTTTACCAGGTGTGGAG<br>GCAGCAGAACGTGCGGGAGTTCCGGCCTTCCTTGAGACGTCGGCGCCGCGCAA<br>TTTACCGTTTTACGAACGTCTTGGATTCACCGTTACGGCGGACGTGGAGGTGCC<br>GGAGGGACCCCGTACTTGGTGTATGACTCGTAAACCGGGAGCCTGATAATCAGT<br>AGCGCTGTCTGGCAACATAAACGGCCCCTTCTGGGCAATGCCGATCAGTTAAGG<br>ATTAGTTGACCGATCCTTAAACTGAGGCACTATAACGGCTTCCACAACAGGGAG<br>CCGTTTTCTTATGCCACTTCTCAATGATCTGCTCGATTTCAGTGACCATCCGCTTA<br>TGCCTCCGCCCTCTGCACAACTATTTGCAGAACACCTTCCCACCGAGTGGATAC<br>AACACTGC |

| | |
|---|---|
| AdhE_F | CTGACAATACGCCTTTTGACAGC |
| AdhE_R | GCGCTCAGGTTCAGACG |
| adhE-conF | GCCGCTGTCTGATAACTGG |
| adhE-conR | GGAAGAGCCATTCCACTGG |
| AdhE-seq1 | CACTATCGCTGAACCAATCGG |
| AdhE-seq2 | AAACTGTCCCCGACTCTG |
| AdhE-seq3 | GCTGCGCTTTATGGATATCCG |

#### 2. Constitutive Promoters

A set of constitutive insulated promoters of varying strength were used for constitutive expression and taken directly from Davis et al., [4] including the proA, proC and proD promoters. A unique terminator was added to the 5' end of constitutive promoters. These were used to drive constitutive pathway expression in growth associated production strains. These promoter sequences are given in Table S1 below.

**Table S5:** Constitutive promoter sequences.

| Promoter | Sequence | Ref. |
|---|---|---|
| BBa_B1004 _proA (GA 1) | CGCCGAAAACCCCGCTTCGGCGGGGTTTTGCCGCACGTCTC CATCGCTTGCCCAAGTTGTGAAGCACAGCTAACACCACGTC GTCCCTATCTGCTGCCCTAGGTCTATGAGTGGTTGCTGGATA ACTTTACGGGCATGCATAAGGCTCGTAGGCTATATTCAGGGA GACCACAACGGTTTCCCTCTACAAATAATTTTGTTTAACTTT | 4 |
| BBa_B1010 _proC (GA 2) | CGCCGCAAACCCCGCCCCTGACAGGGCGGGGTTTCGCCGCA CGTCTCCATCGCTTGCCCAAGTTGTGAAGCACAGCTAACAC CACGTCGTCCCTATCTGCTGCCCTAGGTCTATGAGTGGTTGC TGGATAACTTTACGGGCATGCATAAGGCTCGTATGATATATTC AGGGAGACCACAACGGTTTCCCTCTACAAATAATTTTGTTTA ACTTT | |
| BBa_B1002 _proD (GA 3) | CGCAAAAAACCCCGCTTCGGCGGGGTTTTTTCGCACGTCTC CATCGCTTGCCCAAGTTGTGAAGCACAGCTAACACCACGTC GTCCCTATCTGCTGCCCTAGGTCTATGAGTGGTTGCTGGATA ACTTTACGGGCATGCATAAGGCTCGTATAATATATTCAGGGAG ACCACAACGGTTTCCCTCTACAAATAATTTTGTTTAACTTT | |

##### 3. *Robustness evaluation in Micro-fermentations*

During µL oxygen robustness studies, production culture volume and shaking speed were varied to achieve desired OTR values as previously reported (http://www.enzyscreen.com/oxygen_transfer_rates.htm) [21-22], and as listed below in Table S5. Batch glucose levels during the production stage were altered to assess robustness to glucose. Strains utilized in the robustness experiments at the micro-fermentation scale are listed in Table S6. Results from the micro-fermentation robustness studies are given in Figure S1.

**Table S6:** Culture conditions for different OTR values**.**

| Max OTR | 25mm orbit shaker | | 50mm orbit shaker | |
|---|---|---|---|---|
| | Shaking Speed | Fill Volume | Shaking Speed | Fill Volume |
| (mmol/L-hr) | (rpm) | (µL) | (rpm) | (µL) |
| 25 | 400 | 100 | 300 | 100 |
| 20 | 400 | 150 | 300 | 150 |
| 15 | 400 | 200 | 300 | 200 |

**Table S7:** List of strains used for Micro-fermentation robustness evaluations and their RS1, RS2 and RS3 scores.

| Strain # | Silencing | Proteolysis | Plasmid | RS1 | RS2 | RS3 |
|---|---|---|---|---|---|---|
| 1 | None | None | pSMART-Ala2 | 70.4 | 40.6 | 55.5 |
| 2 | gltA1 | None | pSMART-Ala2 | 68.8 | 45.5 | 57.1 |
| 3 | gltA2 | None | pSMART-Ala2 | 71.7 | 23.9 | 47.8 |
| 4 | g1g2 | None | pSMART-Ala2 | 65.8 | 42.4 | 54.1 |
| 5 | None | G | pSMART-Ala2 | 89.7 | 70.2 | 79.9 |
| 6 | gltA1 | G | pSMART-Ala2 | 45.6 | 23.8 | 34.7 |
| 7 | gltA2 | G | pSMART-Ala2 | 61.8 | 43.9 | 52.8 |
| 8 | g1g2 | G | pSMART-Ala2 | 40.9 | 7.3 | 24.1 |
| 9 | None | F | pSMART-Ala2 | 78.4 | 52.8 | 65.6 |
| 10 | None | U | pSMART-Ala2 | 63.5 | 15.1 | 39.3 |
| 11 | None | FU | pSMART-Ala2 | 81.6 | 66.9 | 74.2 |
| 12 | None | FG | pSMART-Ala2 | 83.0 | 67.9 | 75.4 |
| 13 | gltA1 | FG | pSMART-Ala2 | 66.7 | 49.5 | 58.1 |
| 14 | gltA2 | FG | pSMART-Ala2 | 69.5 | 54.8 | 62.1 |
| 15 | g1g2 | FG | pSMART-Ala2 | 48.5 | 17.0 | 32.8 |
| 16 | None | GU | pSMART-Ala2 | 71.9 | 13.9 | 42.9 |

| | | | | | | |
|---|---|---|---|---|---|---|
| 17 | gltA1 | GU | pSMART-Ala2 | 92.1 | 86.7 | 89.4 |
| 18 | gltA2 | GU | pSMART-Ala2 | 84.0 | 71.9 | 77.9 |
| 19 | g1g2 | GU | pSMART-Ala2 | 93.9 | 83.7 | 88.8 |
| 20 | None | FGU | pSMART-Ala2 | 84.9 | 62.0 | 73.4 |
| 21 | gltA1 | FGU | pSMART-Ala2 | 89.1 | 81.0 | 85.0 |
| 22 | gltA2 | FGU | pSMART-Ala2 | 88.0 | 78.8 | 83.4 |
| 23 | g1g2 | FGU | pSMART-Ala2 | 88.7 | 79.5 | 84.1 |
| 24 | None | None | pSMART-Ala11 | 14.7 | 71.4 | 43.1 |
| 25 | None | None | pSMART-Ala12 | 6.8 | 49.5 | 28.2 |
| 26 | None | None | pSMART-Ala13 | 26.8 | 77.9 | 52.4 |

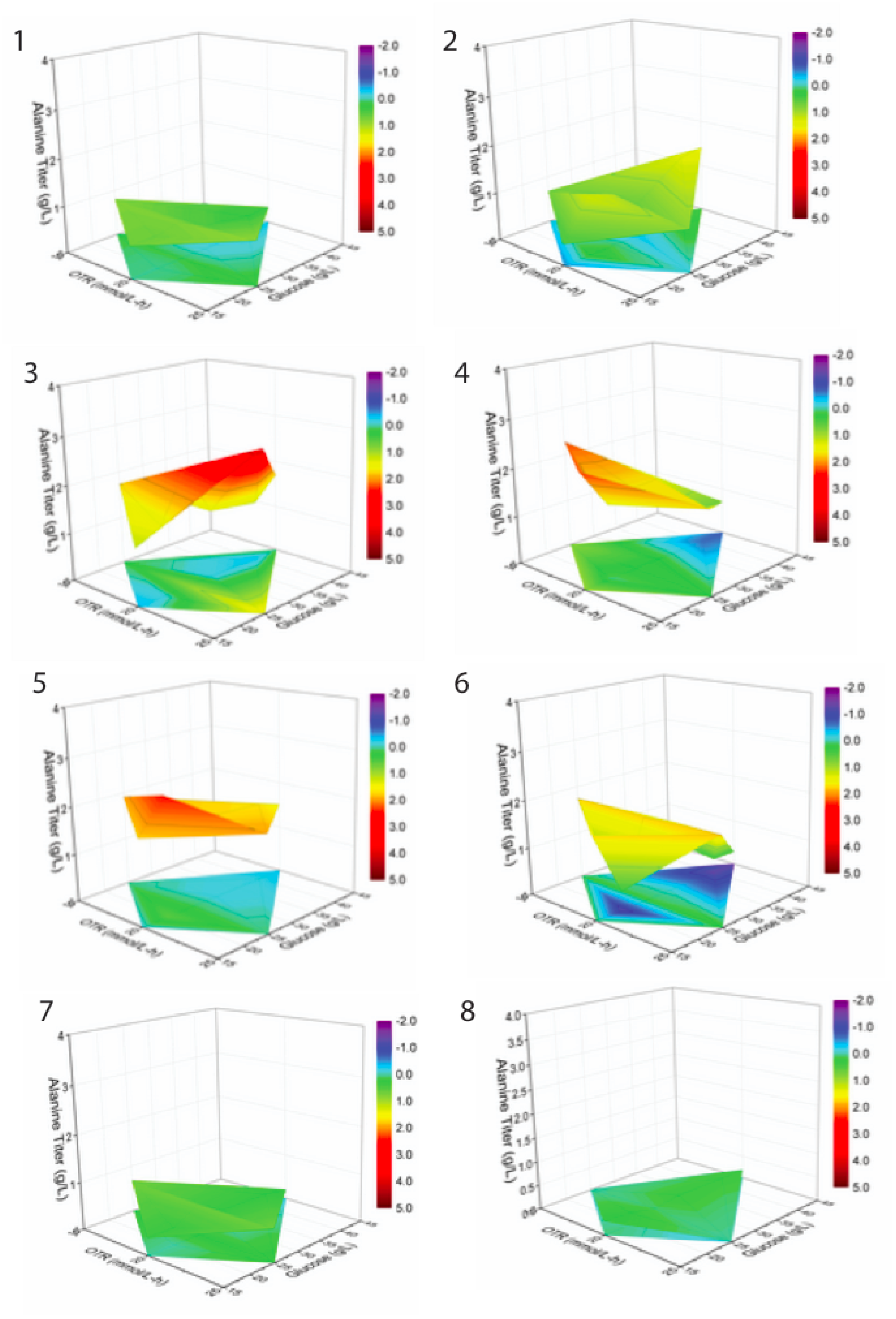
1
2
3
4
5
6
7
8
Alanine Titer (g/L)
OTR (mmol/L-h)
Glucose (g/L)
-2.0
-1.0
0.0
1.0
2.0
3.0
4.0
5.0

9

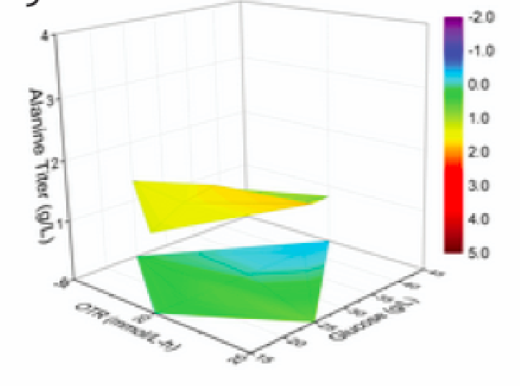


10

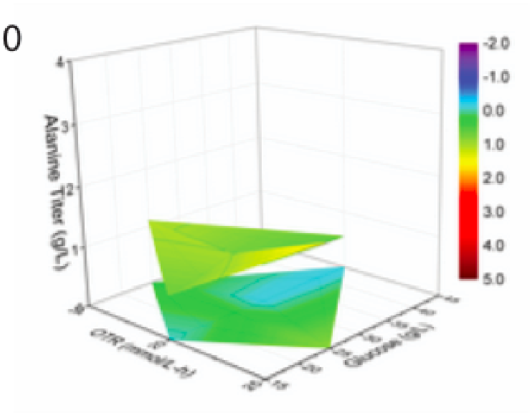


11

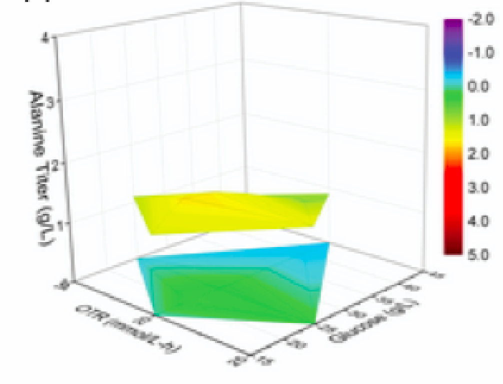


12

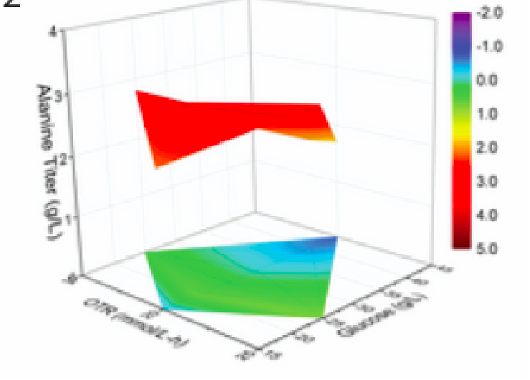


13

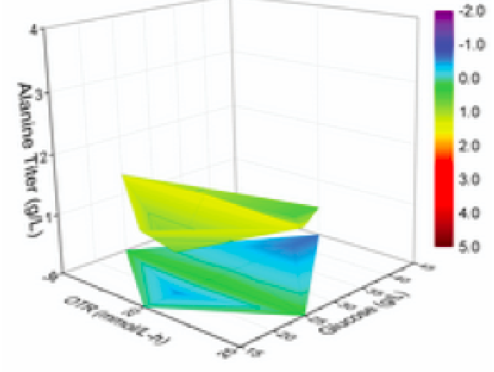


14

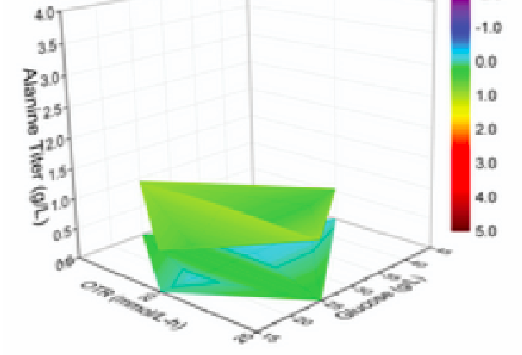


15

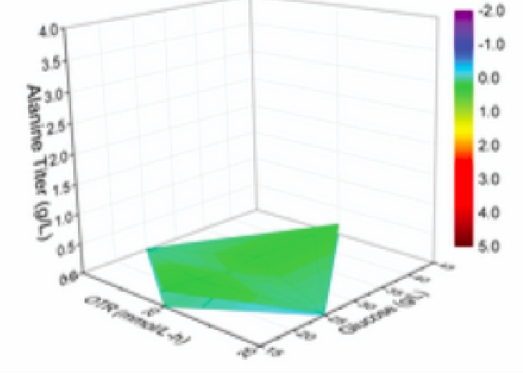


16

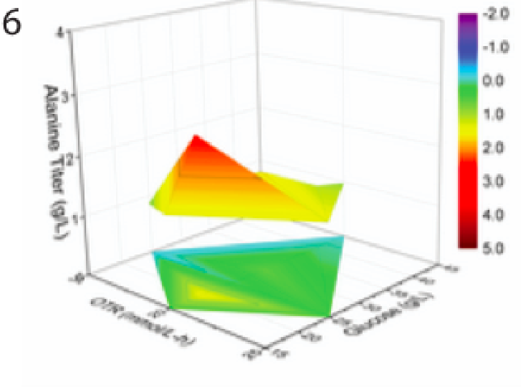

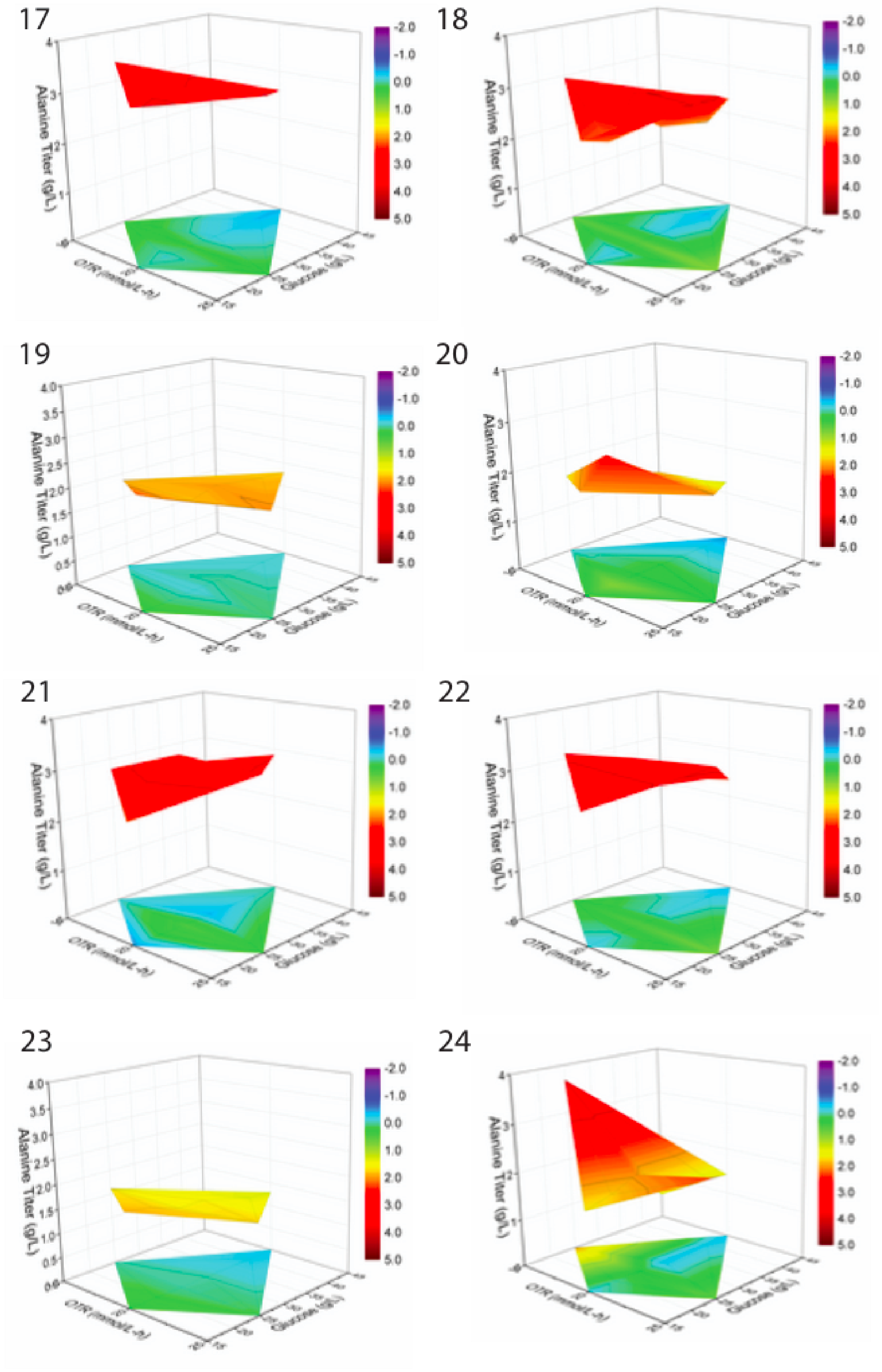
17
18
19
20
21
22
23
24
Alanine Titer (g/L)
OTR (mmol/L-h)
Glucose (g/L)
-2.0
-1.0
0.0
1.0
2.0
3.0
4.0
5.0

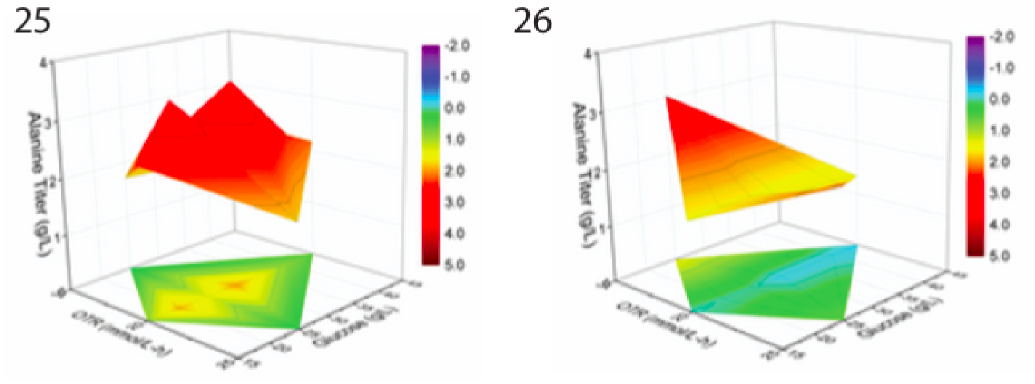


**Figure S1:** Robustness plot for complete strain set. Bottom plane, alanine titer scaled to median, top plane, alanine titer, the same color scale (alanine titer in g/L) was used for both panels. Strain numbering corresponds to Table S6

**Table S8:** Strains used for alanine scalability.

| Strain # | Silencing | Proteolysis | Host Strain | Alanine Plasmid |
|---|---|---|---|---|
| 1 | gltA1 | FGU | DLF_Z0047 | pSMART-Ala2 |
| 2 | G1G2 | G | DLF_Z0043 | pSMART-Ala2 |
| 3 | gltA2 | FG | DLF_Z0039 | pSMART-Ala2 |
| 4 | gltA2 | FGU (+adhE) | DMC_HS_01718 | pSMART-Ala10 |

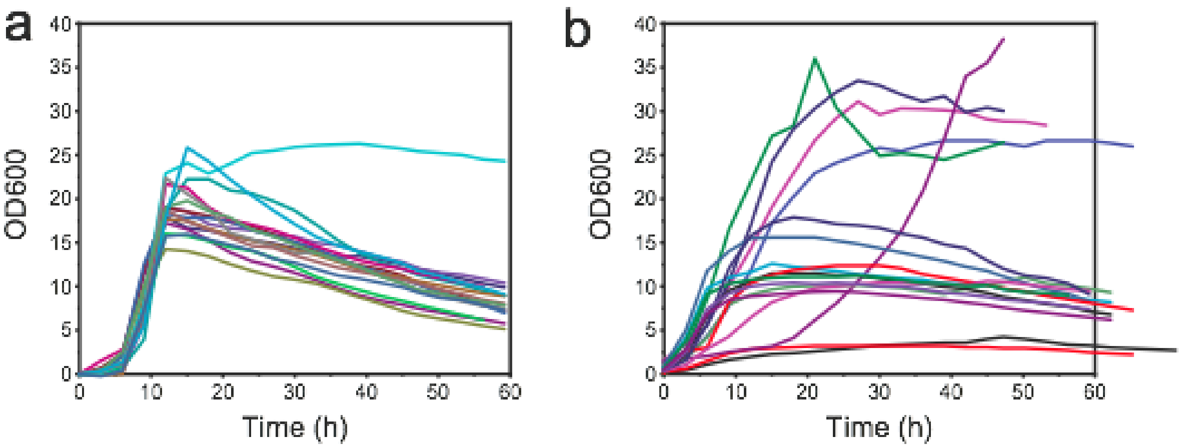


**Figure S2:** Growth profile for all (a) valve and (b) growth associated strains at 1L scale evaluated in this paper**.** Growth curves were synced to account for any variations in lag time. Valve strains growth curves were synced to the same mid-exponential point. Growth associated strains growth curves were synced to the same take-off point.

**Table S9:** Feeding profiles. Feed rate is calculated based on starting volume.

| **Feed Name** | **Feed Start** | **Phase 1 Feeding** | | **Phase 2 Feeding** | | **Phase 3 Feeding** | |
|---|---|---|---|---|---|---|---|
| | | **Feed Rate (g/L-h)** | **Duration (h)** | **Feed Rate (g/L-h)** | **Duration (h)** | **Feed Rate (g/L-h)** | **Duration (h)** |
| 2 g/h* | OD>2 | 2.2 | 50 | | | | |
| 3 g/h* | OD>2 | 3.3 | 50 | | | | |
| 4 g/h* | OD>2 | 4.4 | 50 | | | | |
| Feed_P2 | OD=10~15 | 7.23 | 2.5 | 5.17 | 28.5 | 2.5 | 19 |
| Feed_P12 | OD=10~15 | 10.85 | 2.5 | 7.76 | 28.5 | 3.75 | 19 |

*: used in 1L glucose robustness evaluation.

**Table S10:** Fermentation feeding for all alanine “valve” strains.

| **Scale** | **Strain Information** | | | **Target Biomass (gCDW/L)** | **Feed Name** |
|---|---|---|---|---|---|
| | **Silencing** | **Proteolysis** | **Alanine Plasmid** | | |
| 1L | gltA1 | FGU | pSMART-Ala2 | 10 | 2 g/h, 3 g/h, 4 g/h |
| 1L | G1G2 | G | pSMART-Ala2 | 10 | 2 g/h |
| 1L | gltA2 | FG | pSMART-Ala2 | 10 | 2 g/h |
| 1L | gltA2 | FGU (+adhE) | pSMART-Ala10 | 10 | Feed_P2 |
| 6L | gltA2 | FGU (+adhE) | pSMART-Ala10 | 10 | Feed_P2 |
| 4000L | gltA2 | FGU (+adhE) | pSMART-Ala10 | 10 | Feed_P2 |
| 6L | gltA2 | FGU (+adhE) | pSMART-Ala10 | 20 | Feed_P12 |

**Table S11:** Strains used for citramalate & xylitol scalability in Figure 6.

| Strain | Product | Silencing | Proteolysis | Pathway Plasmid | Source |
|---|---|---|---|---|---|
| 1 | Citramalate | *ev* | None | pHCKan-yibDp-cimA3.7 (Addgene #134595) | [5] |
| 2 | Citramalate | *gltAp1-gltAp2 -zwf* | FGU | pHCKan-yibDp-cimA3.7 (Addgene #134595) | This study |
| 3 | Citramalate | *gltAp2* | G | pHCKan-yibDp-cimA3.7 (Addgene #134595) | [5] |
| 2 | Citramalate | *gltAp2-zwf* | GZ | pHCKan-yibDp-cimA3.7 (Addgene #134595) | [5] |
| 3 | Xylitol | *ev* | None | pHCKan-xyrA (Addgene #58613) | [2] |
| 2 | Xylitol | *gltAp1-zwf* | Z | pHCKan-xyrA (Addgene #58613) | This study |
| 3 | Xylitol | *zwf* | FZ | pHCKan-xyrA (Addgene #58613) | [2] |
| 4 | Xylitol | *zwf* | FZ, pCDF-pntAB | pHCKan-xyrA (Addgene #58613) | [2] |

#### References


1. Li, S., Ye, Z., Lebeau, J., Moreb, E. A. & Lynch, M. D. Dynamic control over feedback regulation improves stationary phase fluxes in engineered E. coli. *bioRxiv* 2020.07.26.219949 (2020) doi:10.1101/2020.07.26.219949.

2. Li, S. *et al.* Dynamic control over feedback regulatory mechanisms improves NADPH fluxes and xylitol biosynthesis in engineered E. coli. *bioRxiv* 2020.07.27.222588 (2020) doi:10.1101/2020.07.27.222588.

3. Menacho‐Melgar, R. *et al.* Scalable, two‐stage, autoinduction of recombinant protein expression in E. coli utilizing phosphate depletion. *Biotechnol. Bioeng.* **26**, 44 (2020).

4. Davis, J. H., Rubin, A. J. & Sauer, R. T. Design, construction and characterization of a set of insulated bacterial promoters. *Nucleic Acids Res.* **39**, 1131–1141 (2011).

5. Shuai Li, Zhixia Ye, Juliana Lebeau Eirik A. Moreb and Michael D. Lynch. Dynamic control over

feedback regulation improves stationary phase fluxes in engineered E. coli. *Materials for Review But not publication*.
